# Integrated plasma proteomic and single-cell immune signaling network signatures demarcate mild, moderate, and severe COVID-19

**DOI:** 10.1101/2021.02.09.430269

**Authors:** Dorien Feyaerts, Julien Hédou, Joshua Gillard, Han Chen, Eileen S. Tsai, Laura S. Peterson, Kazuo Ando, Monali Manohar, Evan Do, Gopal K.R. Dhondalay, Jessica Fitzpatrick, Maja Artandi, Iris Chang, Theo T. Snow, R. Sharon Chinthrajah, Christopher M. Warren, Rich Wittman, Justin G. Meyerowitz, Edward A. Ganio, Ina A. Stelzer, Xiaoyuan Han, Franck Verdonk, Dyani K. Gaudillière, Nilanjan Mukherjee, Amy S. Tsai, Kristen K. Rumer, Sizun Jiang, Sergio Iván Valdés Ferrer, J. Daniel Kelly, David Furman, Nima Aghaeepour, Martin S. Angst, Scott D. Boyd, Benjamin A. Pinsky, Garry P. Nolan, Kari C. Nadeau, Brice Gaudillière, David R. McIlwain

**Affiliations:** Department of Anesthesiology, Perioperative and Pain Medicine, Stanford University School of Medicine, Stanford, CA, USA; Section Pediatric Infectious Diseases, Laboratory of Medical Immunology, Radboud Institute for Molecular Life Sciences, Nijmegen, the Netherlands; Radboud Center for Infectious Diseases, Radboudumc, Nijmegen, the Netherlands; Department of Pathology, Stanford University School of Medicine, Stanford, CA, USA; Department of Microbiology and Immunology, Stanford University School of Medicine, Stanford, CA, USA; Division of Neonatal and Developmental Medicine, Department of Pediatrics, Stanford University School of Medicine, Stanford, CA, USA; Sean N Parker Center for Allergy and Asthma Research, Stanford University, Stanford, CA, USA; Department of Medicine, Stanford University, Stanford, CA, USA; Department of Primary Care and Population Health, Stanford University School of Medicine, Stanford, CA, USA; Division of Allergy, Immunology and Rheumatology, Department of Pediatrics, Stanford University, Stanford, CA, USA; Department of Biomedical Sciences, University of the Pacific, Arthur A. Dugoni School of Dentistry, San Francisco, CA, USA; Division of Plastic & Reconstructive Surgery, Department of Surgery, Stanford University School of Medicine, Stanford, CA, USA; Departamento de Neurología, Instituto Nacional de Ciencias Médicas y Nutrición Salvador Zubirán, Mexico City, Mexico; Department of Epidemiology and Biostatistics, UCSF, San Francisco, CA, USA; Institute of Global Health Sciences, UCSF, San Francisco, CA, USA; F.I. Proctor Foundation, UCSF, San Francisco, CA, USA; Buck Artificial Intelligence Platform, the Buck Institute for Research on Aging, Novato, CA, USA; Stanford 1000 Immunomes Project, Stanford University School of Medicine, Stanford, CA, USA; Austral Institute for Applied Artificial Intelligence, Institute for Research in Translational Medicine (IIMT), Universidad Austral, CONICET, Pilar, Buenos Aires, Argentina; Department of Biomedical Informatics, Stanford University School of Medicine, Stanford, CA, USA; Division of Infectious Diseases and Geographic Medicine, Department of Medicine, Stanford University School of Medicine, Stanford, CA, USA; Department of Medicine, Division of Pulmonary, Allergy and Critical Care Medicine, Stanford University, Stanford, CA, USA

## Abstract

The biological determinants of the wide spectrum of COVID-19 clinical manifestations are not fully understood. Here, over 1400 plasma proteins and 2600 single-cell immune features comprising cell phenotype, basal signaling activity, and signaling responses to inflammatory ligands were assessed in peripheral blood from patients with mild, moderate, and severe COVID-19, at the time of diagnosis. Using an integrated computational approach to analyze the combined plasma and single-cell proteomic data, we identified and independently validated a multivariate model classifying COVID-19 severity (multi-class AUC_training_ = 0.799, p-value = 4.2e-6; multi-class AUC_validation_ = 0.773, p-value = 7.7e-6). Features of this high-dimensional model recapitulated recent COVID-19 related observations of immune perturbations, and revealed novel biological signatures of severity, including the mobilization of elements of the renin-angiotensin system and primary hemostasis, as well as dysregulation of JAK/STAT, MAPK/mTOR, and NF-κB immune signaling networks. These results provide a set of early determinants of COVID-19 severity that may point to therapeutic targets for the prevention of COVID-19 progression.

**Summary:** Feyaerts et al. demonstrate that an integrated analysis of plasma and single-cell proteomics differentiates COVID-19 severity and reveals severity-specific biological signatures associated with the dysregulation of the JAK/STAT, MAPK/mTOR, and NF-κB immune signaling networks and the mobilization of the renin-angiotensin and hemostasis systems.

## Introduction

The ongoing Coronavirus Disease 2019 (COVID-19) pandemic, caused by the novel and highly contagious Severe Acute Respiratory Syndrome Coronavirus-2 (SARS-CoV-2) (Lu et al., 2020b) has affected more than 105 million patients worldwide as of early 2021 (Dong et al., 2020). A wide range of clinical manifestations exists for COVID-19, which require different intervention strategies. While the majority of patients with COVID-19 experience mild or asymptomatic infections, nearly 20% of patients develop severe disease requiring hospitalization (Wu and McGoogan, 2020). A substantial portion (8%-30%) of those hospitalized patients ultimately succumbs to the disease, leading to a devastating global tally of COVID-19 fatalities (CDC, 2021; Horwitz et al., 2021; Piroth et al., 2020; Rosenthal et al., 2020; Wu and McGoogan, 2020).

Varying outcomes for COVID-19 depend on a set of risk factors and the interplay between viral replication, tissue damage, as well as a balance of beneficial and detrimental host immune responses. Several studies have provided evidence for profoundly altered immune responses caused by SARS-CoV-2 infection, including sustained functional changes in circulating immune cells. Lymphopenia (Cao, 2020; Rodriguez et al., 2020), increased inflammatory plasma cytokine levels (Del Valle et al., 2020), dysregulated innate immune cell function (Arunachalam et al., 2020; Schulte-Schrepping et al., 2020; Wilk et al., 2020), and abnormal T cell activation (Mathew et al., 2020) have been observed, particularly in hospitalized patients with severe COVID-19. However, prior studies have primarily focused on patients with severe COVID-19, while fewer studies have included non-hospitalized patients with mild and moderate COVID-19 (Chevrier et al., 2021; Silvin et al., 2020; Su et al., 2020). In addition, while prior studies have reported on the distribution, phenotype, and transcriptional profile of peripheral immune cells, how SARS-CoV-2 infection alters immune cell signaling responses to inflammatory challenges (or immune signaling networks) has not been determined. As such, the immunological mechanisms that differentiate patients with mild, moderate, and severe COVID-19 are poorly understood. Unraveling the underlying immune pathogenesis across the spectrum of COVID-19 presentations is important to both understand the drivers of disease severity as well as to identify clinically relevant biomarkers that can inform therapeutic interventions.

High-dimensional mass cytometry immunoassays are uniquely adapted to the analysis of immune cell signaling networks as multiple intracellular signaling events (e.g. post-translational protein modifications) are simultaneously quantified in precisely phenotyped immune cells, at baseline and in response to *ex vivo* stimulations. The approach has previously enabled the identification of clinically-relevant biological signatures predictive of patient outcomes in several clinical contexts, including infection, malignancies, stroke, and traumatic injury (Ganio et al., 2020; Gaudilliere et al., 2014; Good et al., 2018; Irish and Doxie, 2014; Myklebust et al., 2017; Rahil et al., 2020; Tsai et al., 2019).

In this study, we combined the mass-cytometry analysis of immune cell signaling responses with the high-content proteomic analysis of plasma analytes in blood samples from patients with mild, moderate, and severe COVID-19 to identify biological signatures that demarcate COVID-19 clinical manifestations. The integrated single-cell and plasma proteomic analysis allowed including an additional dimension in the characterization of immune signaling networks by accounting for the plasma environment of circulating immune cells.

## Results

### Combined plasma and single-cell proteomic analysis of peripheral blood samples from patients with mild, moderate, and severe COVID-19

97 SARS-CoV-2 positive patients with mild, moderate or severe COVID-19 were enrolled in this study at Stanford University Medical Center in California, USA (**Figure 1A**). Patient characteristics can be found in **Table 1**. COVID-19 severity was determined using previously defined clinical criteria ((Chen et al., 2020); see Methods). For each patient a peripheral blood sample was collected at the time of COVID-19 diagnosis. Samples from SARS-CoV-2 positive patients were examined alongside those from 40 healthy controls collected at Stanford in 2019, before the detection of SARS-CoV-2 in the geographic region. Blood samples were used to isolate both plasma and peripheral blood mononuclear cells (PBMC). PBMC were stimulated *ex vivo* to trigger pathogen sensing and cytokine signaling response pathways in innate and adaptive immune cells relevant during infection (TLR4 stimulant LPS and TLR7/8 agonist CL097; IFN*α*, IL-2, IL-4, and IL-6 cytokine cocktail; and cell stimulation cocktail consisting of phorbol 12-myristate 13-acetate (PMA) and Ionomycin (I) (PI)) (**Figure 1B**). The frequencies of 44 manually-gated immune cell subsets representing major circulating innate and adaptive immune cells were determined using a 42-parameter single-cell phospho-mass cytometry immunoassay (**Figure S1**). For each immune cell subset, the frequency was determined, alongside baseline signaling activity (unstimulated condition) and signaling response capacity of cells to *ex vivo* stimulation with inflammatory reagents, which were measured by examining the phosphorylation state of 15 intracellular signaling proteins. The mass cytometry analysis resulted in a total of 2662 immune cell response features per PBMC sample. To complement this single-cell analysis, 1472 circulating plasma proteins were measured using the proximity extension assay (PEA) platform from Olink Proteomics. Five mass cytometry (cell frequency, baseline signaling, IFN*α*/IL-2/IL-4/IL-6 signaling response, LPS/CL097 signaling response, and PI signaling response) and one plasma proteomic data layer(s) were collected, resulting in six data layers in total. Correlation networks demonstrate the existence of strong inter-and intra-layer correlations between features (**Figure 1C-D**).

**Figure 1.**
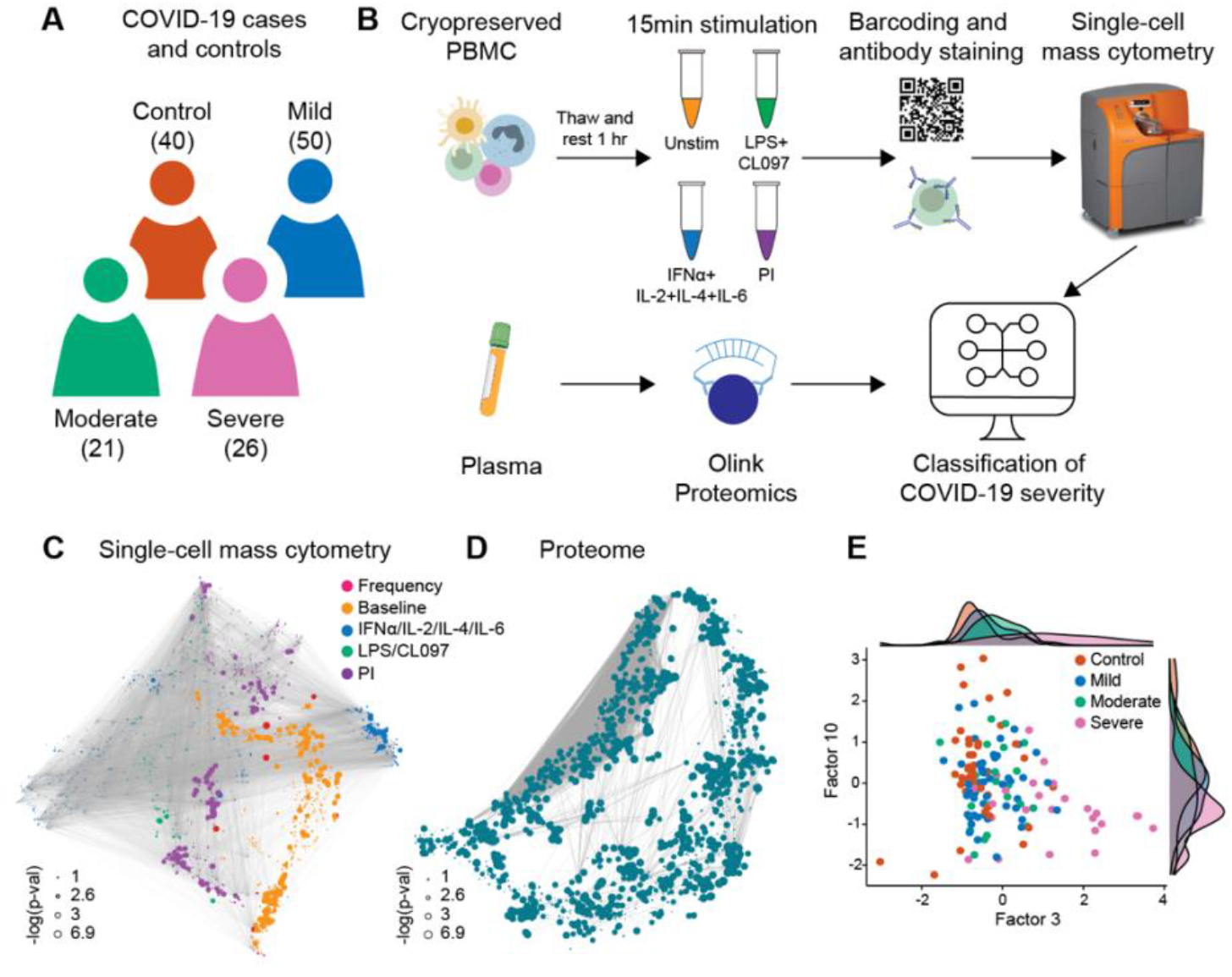
Combined plasma and single-cell proteomic profiling of patients with mild, moderate, and severe COVID-19. **A** Patients with mild (50), moderate (21), and severe (26) COVID-19 were examined together with 40 healthy controls. **B** Schematic representation of the experimental workflow. Plasma proteins were measured using the Olink Explore 1536 assay, while PBMC were stimulated with either LPS+CL097, IFN*α*+IL-2+IL-4+IL-6, PMA+ionomycin (PI), or left unstimulated (Unstim) before barcoding, antibody staining, and analysis by single-cell mass cytometry. **C-D** Correlation networks of single-cell mass cytometry and proteome dataset. Each node represents a feature, with edges representing the correlation between features (cor > 0.9). Node size reflects *-log_10_* of p-value of the correlation with severity (Spearman) and node color represents the different data layers. **E** Bivariate scatterplot of patients with COVID-19 and healthy controls plotted along factors 3 and 10 identified by multi-omic factor analysis (MOFA; see Figure S2).

**Table 1.** Patient characteristics. Data are shown as a number and a percentage. Age is reported in median years (minimum to maximum).

|  | **Total (N =137)** |  | **Control (N=40)** |  | **COVID+ (N=97)** |  |
| --- | --- | --- | --- | --- | --- | --- |
| **Age** | 45 (19-78) |  | 48 (23-73) |  | 44 (19-78) |  |
| **Sex** | **N** | **%** | **N** | **%** | **N** | **%** |
| M | 59 | 43.07 | 16 | 40 | 43 | 44.33 |
| F | 78 | 56.93 | 24 | 60 | 54 | 55.67 |
| **Ethnicity** | **N** | **%** | **N** | **%** | **N** | **%** |
| Asian | 34 | 24.82 | 11 | 27.5 | 23 | 23.71 |
| Hispanic/Latino | 33 | 24.09 | 0 | 0 | 33 | 34.02 |
| Caucasian | 49 | 35.77 | 15 | 37.5 | 34 | 35.05 |
| Black | 6 | 4.38 | 1 | 2.5 | 5 | 5.15 |
| Multiracial | 5 | 3.65 | 3 | 7.5 | 2 | 2.06 |
| Middle Eastern | 1 | 0.73 | 1 | 2.5 | 0 | 0 |
| Not reported | 9 | 6.57 | 9 | 22.5 | 0 | 0 |
| **Disease Status** | **N** | **%** | **N** | **%** | **N** | **%** |
| Healthy | 40 | 29.2 | 40 | 100 | 0 | 0 |
| Asymptomatic | 2 | 1.46 | 0 | 0 | 2 | 2.06 |
| Mild | 48 | 35.04 | 0 | 0 | 48 | 49.48 |
| Moderate | 21 | 15.33 | 0 | 0 | 21 | 21.65 |
| Severe | 26 | 18.98 | 0 | 0 | 26 | 26.8 |
| **Comorbidities** |  |  |  |  |  |  |
| **Diabetes** | **N** | **%** | **N** | **%** | **N** | **%** |
| Yes | 13 | 9.49 | 0 | 0 | 13 | 13.4 |
| Prediabetic | 5 | 3.65 | 1 | 2.5 | 4 | 4.12 |
| No | 96 | 70.07 | 32 | 80 | 64 | 65.98 |
| Not reported | 23 | 16.79 | 7 | 17.5 | 16 | 16.5 |
| **Asthma** | **N** | **%** | **N** | **%** | **N** | **%** |
| Yes | 24 | 17.52 | 4 | 10 | 20 | 20.62 |
| No | 85 | 62.04 | 29 | 72.5 | 56 | 57.73 |
| Not reported | 28 | 20.44 | 7 | 17.5 | 21 | 21.65 |
| **CV Condition** | **N** | **%** | **N** | **%** | **N** | **%** |
| Yes | 15 | 10.95 | 5 | 12.5 | 10 | 10.31 |
| No | 115 | 83.94 | 28 | 70 | 87 | 89.69 |
| Not reported | 7 | 5.11 | 7 | 17.5 | 0 | 0 |
| **Obesity** | **N** | **%** | **N** | **%** | **N** | **%** |
| Yes | 18 | 13.14 | 0 | 0 | 18 | 18.6 |
| No | 89 | 64.96 | 33 | 82.5 | 56 | 57.7 |
| Not reported | 30 | 21.9 | 7 | 17.5 | 23 | 23.7 |

To understand the relationships between the different data layers and COVID-19 severity, we first applied unsupervised dimensionality reduction with multi-omics factor analysis (MOFA; (Argelaguet et al., 2018)), which infers a set of factors that capture shared sources of variability across different datasets. We supplied all six data layers for the analysis, resulting in a MOFA model with 17 factors (**Figure S2A**). Both single-cell immune response (baseline signaling and signaling response to IFN*α*/IL-2/IL-4/IL-6 and LPS/CL097; factor 10) and plasma proteome (factor 3) data contributed strongly to the variance observed in our samples (**Figure S2B**) and were significantly associated with COVID-19 severity (**Figure S2C**). A gradient with increasing disease severity across factors 3 and 10 (**Figure 1E**) was also observed, suggesting that single-cell immune response and plasma proteome data both contain clinically important biological events. The results prompted us to perform an integrated analysis to determine whether a classifier of COVID-19 severity could be derived from the combined plasma and single-cell proteomics data.

### Integrated modeling of plasma and single-cell proteome differentiates COVID-19 severity

A high-dimensional computational analysis pipeline was applied to train and independently validate (training cohort: n=74; 25 control, 20 mild, 10 moderate, and 19 severe patients; validation cohort: n=63; 15 control, 30 mild, 11 moderate, and 7 severe patients) an integrated model of COVID-19 severity based on the combined proteomic and single-cell immune response data (Ghaemi et al., 2019). In this approach, the six data layers were considered separately and a two-step process was used to combine these data layers in a multi-omic fashion (**Figure 2A**). Cross-validated multivariate Least Absolute Shrinkage and Selection Operator (LASSO) linear regression models (Tibshirani, 1996) were first trained for each individual data layer of the training cohort, with disease severity used as a ranked order variable (i.e., control classed as 1 to severe classed as 4), and second, the individual prediction models were integrated into a single model by stacked generalization (SG) (Ghaemi et al., 2019) (**Figure 2A**). The second step uses the estimations of disease severity of each LASSO model as predictors for a constrained regression model.

**Figure 2.**
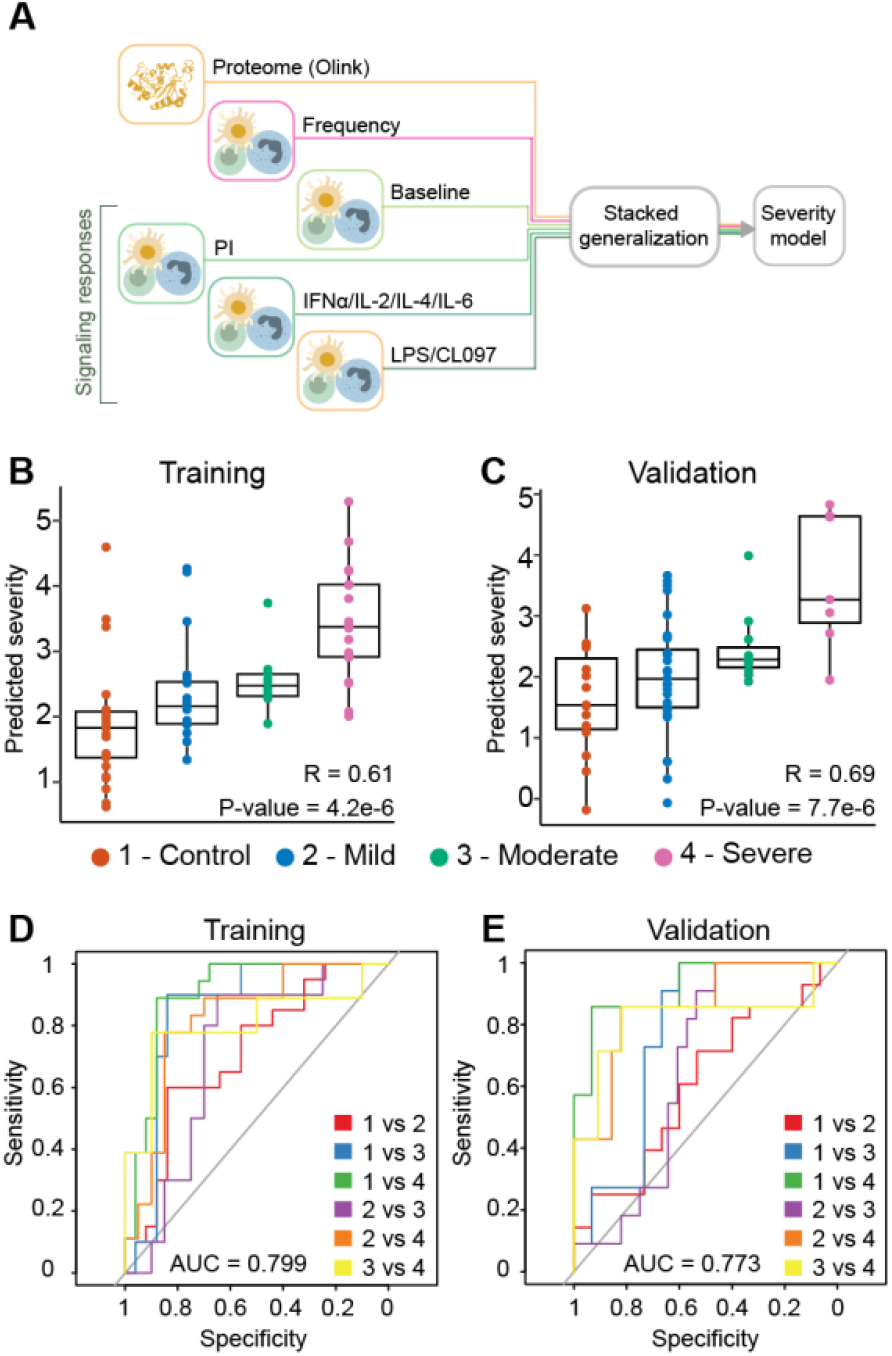
Integrated modeling of plasma and single-cell proteomic events categorizes COVID-19 severity. **A** Integration of all six data layers (proteome, frequency, baseline signaling, LPS/CL097 signaling response, IFN*α*/IL-2/IL-4/IL-6 signaling response, and PI signaling response) using a stacked generalization (SG) method. **B-C** Outcome of predicted vs true disease severity derived from SG model for the **B** training (R = 0.61, p-value = 4.2e-6, n = 74) and **C** validation cohort (R = 0.69, p-value = 7.7e-6, n = 73). **D-E** Multi-class area under the curve receiver operating characteristic (ROC) analysis of **D** the training (AUC = 0.799, n = 74) and **E** validation (AUC = 0.773, n = 73) severity model (see Table S2 for individual AUCs). 1 = control, 2 = mild, 3 = moderate, and 4 = severe.

The analysis identified a predictive SG model (“severity model”) that classified COVID-19 severity categories at time of sampling for patients in the training cohort (R = 0.61, p-value = 4.2e-6, n=74). The generalizability of the severity model was independently tested in patients from the validation cohort (R = 0.69, p-value = 7.7e-6, n=63) (**Figure 2B-C**). The contribution of individual data layers to the overall severity model was highest for the plasma proteome and lowest for the signaling responses to LPS/CL097 stimulation, according to severity model coefficients (**Table S1**).

To estimate the performance of the severity model, a multi-class area under the curve receiver operating characteristic (ROC) analysis was performed for the training and validation cohort (**Figure 2D-E**). The multi-class ROC analysis showed that the severity model performed well at classifying patients across disease severity categories (multi-class AUC_training_ = 0.799; multi-class AUC_validation_ = 0.773). The most accurate classification was achieved when classifying severe patients from the other patient groups (**Table S2**).

Results from the severity model indicate that mild, moderate, and severe COVID-19 manifestations can be differentiated from the measurement of plasma proteins and single-cell immune signaling events in patient’s peripheral blood. A confounder analysis further showed that the severity model remained significant when accounting for clinical or demographic variables that have previously been associated with COVID-19 severity, including age, sex, BMI, and time between symptom onset and blood collection (**Table S3**).

### Biological signatures of COVID-19 severity

To facilitate the biological interpretation of the high-dimensional severity model and to identify potential biomarker targets, the contribution of individual plasma and single-cell proteomic features to the severity model performance was quantified by measuring repeated selection during a 1,000-iteration bootstrap procedure (**Figure 3A**; (Bach, 2008; Chatterjee and Lahiri, 2011)). We examined in detail the set of features that were within the top 10% selected by the bootstrap procedure, and that thus contributed strongly to the overall model (**Figure 3B-C and supplemental data file S1)**. Single-cell (**Figure 3B**) and plasma (**Figure 3C**) proteomic features were visualized with two correlation networks, highlighting the interconnectivity of the proteomic datasets (edges), and the correlation between individual features and COVID-19 severity (node size/color). Features within the top 10% of bootstrap selection segregated into correlated communities. Single-cell immune response features were annotated according to the cellular attribute (immune cell frequency or signaling response) most commonly represented within each community (**Figure 3B**), while plasma proteomic features were annotated by protein name (**Figure 3C**). These communities of informative features are discussed below in relation to disease severity.

**Figure 3.**
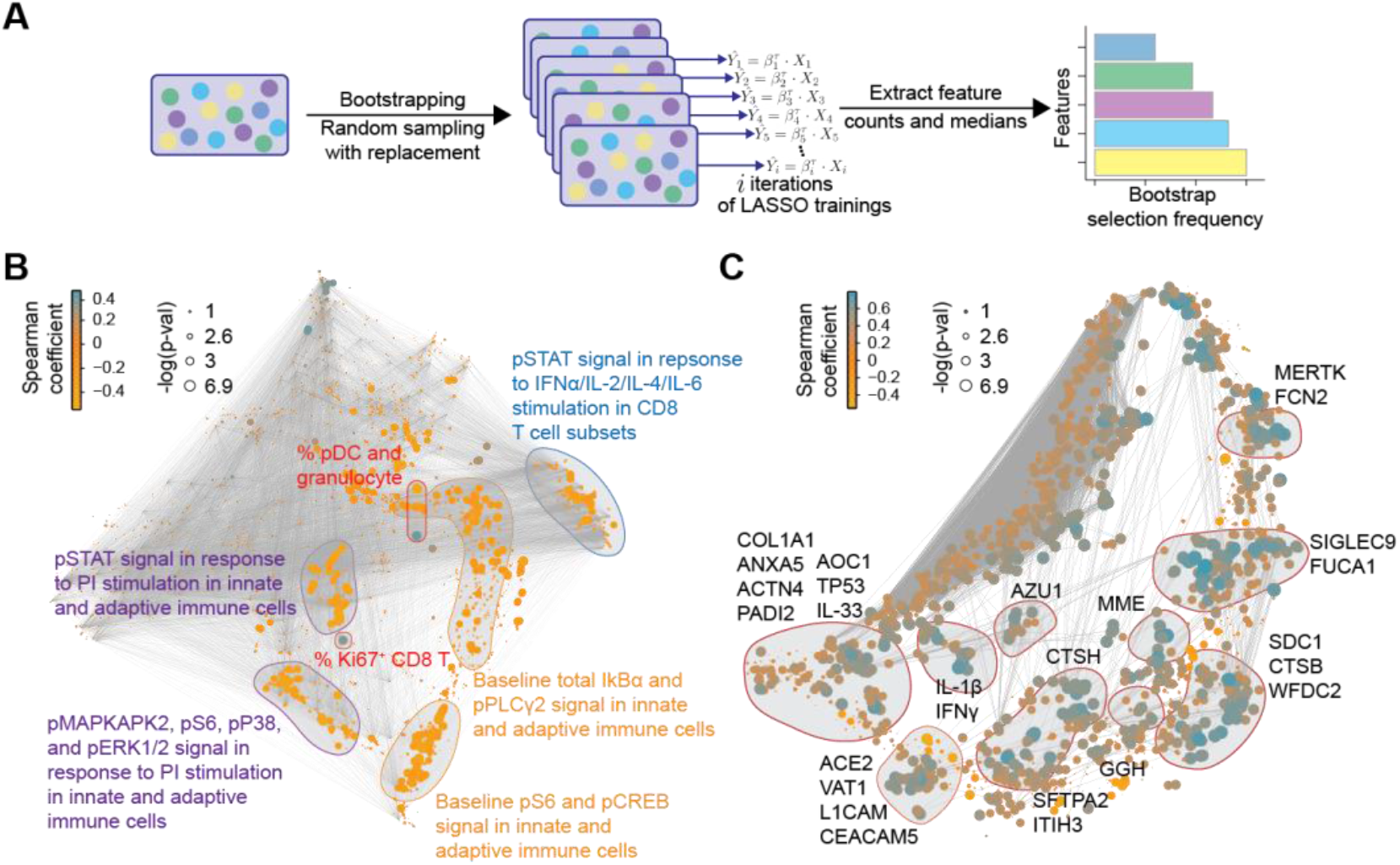
An iterative bootstrapping method identifies robust informative features for the differentiation of mild, moderate, and severe COVID-19. **A** Workflow of the iterative bootstrap method used to identify informative features in the six data layers of the severity model. A LASSO regression model was run 1,000 times on random sub-samples with replacement for each data layer, Xi, then the number of times an individual feature was selected in one of the bootstrap iterations was counted and the features were ranked according to the frequency of selection in the bootstrap models. **B-C** Correlation network depicting single-cell (**B**) or plasma (**C**) proteomic features. Edges represent the correlation between features (Spearman cor > 0.9). Blue/orange nodes highlight positive/negative correlation with disease severity. Node size reflects *-log_10_* of p-value (Spearman). Communities containing the bootstrap-selected informative single-cell (**B**) or plasma (**C**) proteomic features are highlighted and annotated.

Cell frequency features prominent in the correlation network and contributing to the severity model revealed changes in immune cell distribution that are reminiscent of recent immunophenotyping studies in patients with severe COVID-19. For instance, in our study and prior reports, plasmacytoid dendritic cells (pDC) and CD161^+^CD8^+^ T cell frequencies were negatively correlated, while Ki67^+^CD8^+^ T cell and granulocyte frequencies were positively correlated with COVID-19 severity (Arunachalam et al., 2020; Chevrier et al., 2021; Kuri-Cervantes et al., 2020; Mathew et al., 2020; Parrot et al., 2020) (**Figure 4A**). Furthermore, a positive correlation of plasmablast frequency with severity was evident in our data, complementing prior reports (Kuri-Cervantes et al., 2020; Mathew et al., 2020) (**Figure S3A**).

**Figure 4.**
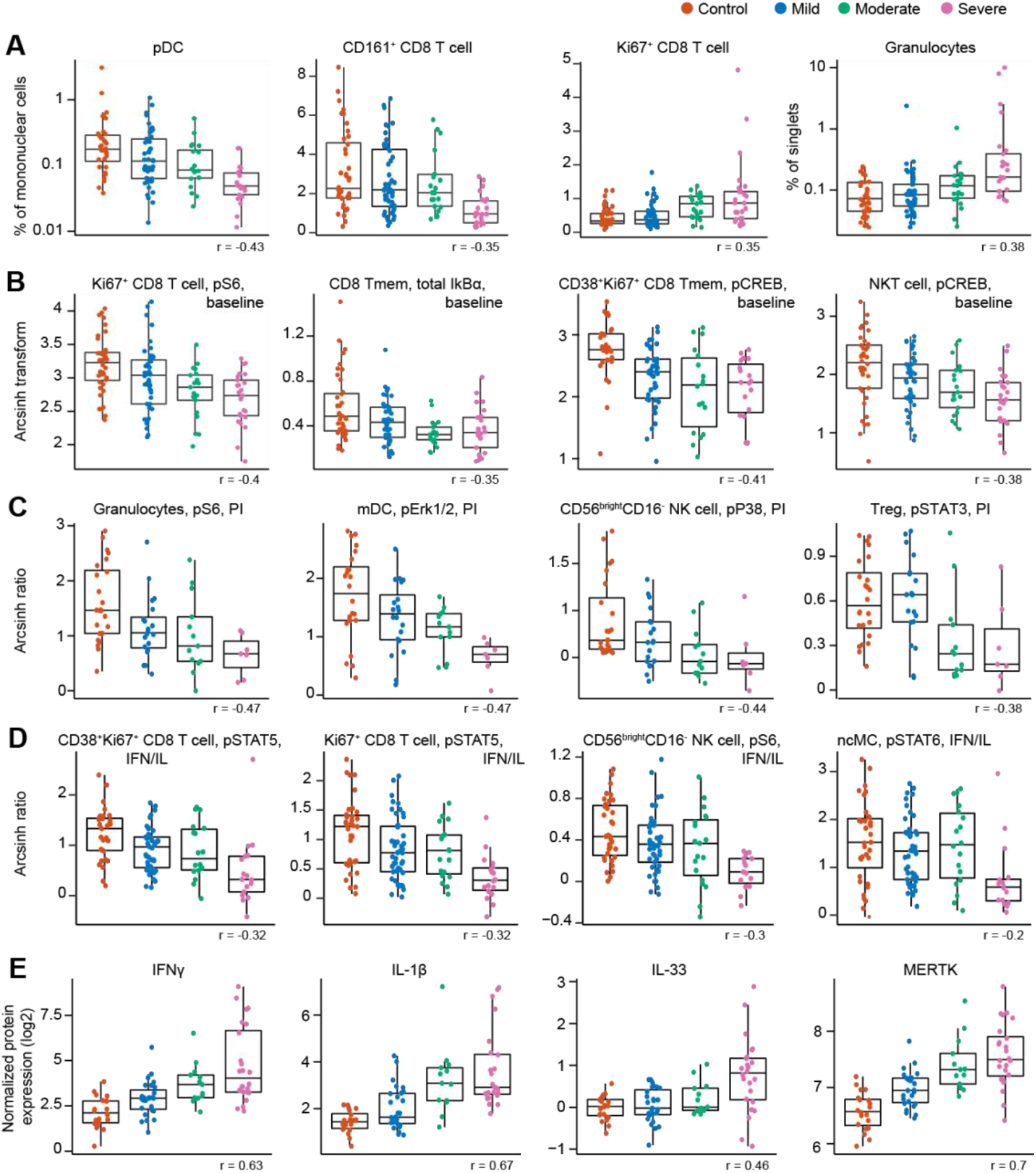
Severity model features reveal biological signatures that demarcate patients with mild, moderate, and severe COVID-19. Boxplots, classified by disease severity, showing features informative to the model, including the correlation of the feature with disease severity (Spearman coefficient). **A** Frequency features are shown as a percentage of mononuclear cells. Granulocytes are shown as a percentage of singlets. pDC and granulocyte frequency are plotted on a log-scale. **B** Immune cell signaling at baseline (arcsinh transformed values; see methods). **C** Immune cell signaling response to PI stimulation is reported as the arcsinh transformed ratio over the baseline signaling response (see methods). **D** Immune cell signaling responses to IFN*α*/IL-2/IL-4/IL-6 (IFN/IL) stimulation are reported as the arcsinh transformed ratio over the baseline signaling response (see methods). **E** Plasma protein levels are reported as the normalized protein expression, an arbitrary unit provided by the Olink assay. Tmem = memory T cell; MERTK = tyrosine-protein kinase Mer.

Assays examining baseline and stimulation dependent cell signaling markers revealed alterations in cell states associated with varying disease severity. A negative correlation was observed between baseline signaling in CD8^+^ T cell subsets and NKT cells, and COVID-19 severity, notably for the phosphorylated (p) S6, total IkB*α*, and pCREB signals (**Figure 4B and S3B**). Negative correlations with COVID-19 severity were also observed for the p4EBP1 and total IkB*α* signals at baseline in granulocytes, while granulocyte pERK1/2 signal at baseline was positively correlated with disease severity (**Figure S3B**). Overall, this data suggests that alterations in immune cell signaling markers at baseline exist between COVID-19 severity categories. In particular, baseline signaling activities of key elements of the mTOR, MAPK, and NF-κB pathway (pS6, pERK1/2, pCREB, and total IkB*α*) were diminished with increasing disease severity in immune cell populations that play important roles in the defense against viral pathogens, such as CD8^+^ T, NKT, and granulocyte cell subsets.

Additionally, immune cell signaling responses to PI stimulation were largely diminished with increasing disease severity, especially for innate immune cell populations. Specifically, PI-stimulated signaling responses in granulocytes (pS6, pERK1/2, and pP38), mDCs (pERK1/2 and pSTAT3), and CD56^bright^CD16^-^ NK cells (pP38) were negatively correlated with disease severity (**Figure 4C and S3C**). Negative correlations with disease severity were also observed for the pSTAT3 signal in Treg and pMAPKAPK2 signal in CD161^+^ CD8^+^ T cells in response to PI stimulation (**Figure 4C and S3C**). Decreased responsiveness to PI stimulation may be an indication of reduced effector function of circulating innate and adaptive immune cells during severe COVID-19, driven by cell-intrinsic effects and/or changes in cytokines and other modulating factors present in the circulation.

For immune cell signaling responses to IFN*α*/IL-2/IL-4/IL-6 stimulation, we also observed a negative correlation with disease severity for the pSTAT4/5/6 signals in both adaptive and innate immune cells (**Figure 4D and S3D**), suggestive of an impaired signaling response to interferon and cytokines in those individuals with more severe COVID-19. Indeed, others have also observed an impaired type I interferon activity in peripheral immune cells of patients critically ill with COVID-19, shown by downregulation of interferon-stimulated genes upon whole blood IFN*α* stimulation (Hadjadj et al., 2020).

Among the most robust plasma proteomic features used by the severity model were several features that positively correlated with disease severity and which overlapped with prior descriptions of the cytokine storm syndrome described in patients with severe COVID-19 (Cao, 2020). For instance, plasma levels of the cytokines IFNγ, IL-1β, and IL-33, showed a positive correlation with disease severity (**Figure 4E**). IL-6, one of the first plasma cytokines recognized as elevated during COVID-19 (Ruan et al., 2020), also exhibited a positive correlation with disease severity in our cohort (**Figure S4**). Although this feature contributed to the severity model, it ranked well below the top 10% of bootstrap selected informative features (see **supplemental data file S1**). A similar positive correlation was observed for lung-related proteins in circulation, such as pulmonary surfactant-associated protein A2 (SFTPA2) and Cathepsin H (CTSH), which are involved in surfactant homeostasis (**Figure S4**) (Buhling et al., 2011; Thorenoor et al., 2018).

Angiotensin-converting enzyme 2 (ACE2) is used as a viral entry receptor by SARS-CoV-2 and is released from the epithelial cell surface upon viral binding (Mortaz et al., 2020). ACE2 ranked within the top 10% of informative features, and levels showed a positive correlation with severity (**Figure S4**). The severity model also identified the protease MME (neprilysin), another key player of the Renin-Angiotensin System (RAS) (Chappell, 2019), as a positive correlate of COVID-19 severity (**Figure S4**). MME is also implicated in neutrophil degranulation (Didangelos, 2020) and our list of informative model features contained multiple other proteins involved in neutrophil degranulation (protein pathway identified by Reactome, see methods; Reactome gene set identifier R-HSA-6798695.2; **Figure S5A**), which were mainly positively correlated with severity as well. This dysregulation of neutrophil degranulation in severe patients is in agreement with recent plasma proteome findings (Overmyer et al., 2020).

Our analysis revealed tyrosine-protein kinase Mer (MERTK) as the most robust feature in the proteome dataset contributing to the severity model (**supplemental data file S1**). Levels of MERTK, an immunosuppressive tyrosine kinase receptor (Lemke and Silverman, 2020), were positively correlated with disease severity (**Figure 4E**). MERTK is found on the surface of macrophages where it mediates phagocytosis of apoptotic cells (Cai et al., 2018). Activation of MERTK has an immunosuppressive effect by downregulating the production of cytokines and type I interferons (Cai et al., 2018; Lemke and Silverman, 2020). Increased MERTK shedding could result in reduced surface expression and loss of MERTK signaling (Sather et al., 2007), which could play a central role in the hyper inflammation observed in severe COVID-19. MERTK also plays a role in platelet aggregation (Sather et al., 2007) and endothelial barrier integrity (Li et al., 2019).

Plasma levels of several other proteins involved in primary hemostasis (Reactome gene set identifier R-HSA-109582) were found to be informative in our model of disease severity (**Figure S5B**), with the majority of them displaying positive correlations with disease severity.

In summary, informative features of the severity model revealed biological signatures that progressed from mild to moderate and severe COVID-19. Salient characteristics of this biological progression included cellular elements of immune signaling networks implicated in defensive immunity against viral pathogens (such as the progressive dampening of NF-κB, MAPK/mTOR, and JAK/STAT signaling in multiple innate and adaptive immune cell subsets) and sentinel proteomic pathways involved in lung and RAS homeostasis, primary hemostasis, neutrophil degranulation, and inflammation.

## Discussion

This study combined high-content plasma proteomics with the single-cell analysis of immune signaling responses to identify biological determinants of severity across the spectrum of COVID-19 manifestations. Using a two-step analytical approach that accounts for the dimensionality of different data layers, we built and independently validated an integrated model that classifies COVID-19 severity. The biological underpinnings of the severity model consisted of co-regulated plasma and single-cell proteomic elements that progressed with COVID-19 severity, including the inflammatory cytokine response to SARS-CoV-2, the mobilization of the Renin-Angiotensin (RAS) and primary hemostasis systems, and the dysregulation of the JAK/STAT, NF-κB and MAPK/mTOR immune signaling responses. The identification of biological signatures progressing with COVID-19 severity provides a set of sentinel events detectable in the early phase of infection that may be targeted therapeutically for the prevention of severe COVID-19.

The ongoing pandemic has fueled major research efforts towards understanding the host immune response against SARS-CoV-2 infection (Arunachalam et al., 2020; Cao, 2020; Del Valle et al., 2020; Mathew et al., 2020; Schulte-Schrepping et al., 2020; Wilk et al., 2020). Previous efforts have been particularly focused on hospitalized patients with severe COVID-19, which, while of paramount importance, excludes the majority of SARS-CoV-2-infected patients who suffer from mild or moderate COVID-19 and do not require hospitalization. The comparative analysis of samples from patients with mild, moderate, and severe disease afforded a more exhaustive characterization of immune responses related to COVID-19 severity. Our approach dovetails with prior studies identifying an immunological shift distinguishing the spectrum of COVID-19 infection states (Chevrier et al., 2021; Su et al., 2020). Consistent with our results, this switch included an increase in inflammation, the emergence of CD4 and CD8 T cells with a proliferative-exhausted phenotype, and a distinct activated myeloid signature (Chevrier et al., 2021; Su et al., 2020). In previous work, immune cell function and responses are indirectly inferred through either phenotype or baseline single-cell mRNA transcriptomics changes, while our approach provided a direct assessment of baseline intracellular signaling responses of multiple immune cell subsets as well as their capacity to respond to inflammatory stimulation.

Two major biological signatures associated with the progression from mild to moderate and severe disease emerged from our integrated analysis: 1) the dampening of NF-κB, MAPK/mTOR and JAK/STAT intracellular signaling responses in multiple innate and adaptive immune cell subsets; and 2) the mobilization of a proteomic network enriched for elements of the RAS, lung homeostasis and hemostasis pathways, alongside canonical elements of the cytokine storm signature of severe COVID-19.

Several of the features informative to our severity model resonate with previous findings in patients suffering from severe COVID-19. For example, plasma levels of cytokines such as IFNγ, IL-1β, IL-33, and IL-6 increased with increasing severity, consistent with the cytokine storm observed in severe patients with COVID-19 (Cao, 2020), and with circulating IL-33 levels as a potential indicator of damaged lung tissue (Liew et al., 2010; Martin and Martin, 2016; Zizzo and Cohen, 2020). In addition, changing frequencies of circulating immune cells also aligned with prior reports in severe patients (Arunachalam et al., 2020; Kuri-Cervantes et al., 2020; Mathew et al., 2020; Parrot et al., 2020; Wilk et al., 2020). Interestingly, an increased percentage of granulocytes was observed in the PBMC fraction of patients with severe COVID-19 (Chevrier et al., 2021; Schulte-Schrepping et al., 2020; Wilk et al., 2020). Low-density granulocytes also appear in the PBMC fraction of patients with inflammatory diseases and severe infection (Hassani et al., 2020; Silvestre-Roig et al., 2019).

The functional analysis of intracellular signaling events in this study revealed intriguing new biology associated with COVID-19 severity, notably with respect to immune cell responses to inflammatory ligands. The aforementioned low-density granulocytes can display dysfunctional immune responses (Hassani et al., 2020; Silvestre-Roig et al., 2019), which dovetails with our observation of decreased immune cell signaling response by the granulocytes present in the PBMC fraction of patients with severe disease. In addition, in agreement with Overmyer *et al*. (Overmyer et al., 2020), increasing plasma levels of several proteins involved in neutrophil degranulation were correlated with disease severity. Excessive release of granules can result in tissue damage and is a feature of acute lung injury and septic shock (Lacy, 2006). In addition to granulocytes, other cell types such as mDCs, NK cells, NKT cells, Tregs, and CD4 and CD8 T cells also showed an inverse relationship between capacity to respond to cytokine stimulation and disease severity, suggestive of overall diminished effector functions of circulating innate and adaptive immune cells with increasing severity. Indeed, others have shown decreased functional responses and exhausted phenotypes in peripheral innate and adaptive immune cells in severe patients as well (Arunachalam et al., 2020; Hadjadj et al., 2020; Kusnadi et al., 2021; Remy et al., 2020b; Wilk et al., 2020). As such, restoring the effector responses of circulating immune cells by immune potentiators (e.g. immune checkpoint inhibitors) to enhance host immunity while simultaneously controlling the cytokine storm (e.g. corticosteroids) may be beneficial in preventing severe disease and overcoming infection (Florindo et al., 2020; Remy et al., 2020a; Tang et al., 2020).

In addition to dysregulated immune signaling responses, the examination of the severity model features revealed several key proteomic pathways that were mobilized with disease severity, including pathways related to lung homeostasis, RAS homeostasis, and hemostasis. Notably the plasma levels of two proteins implicated in the production of lung surfactant (SFTPA2 and CTSH; (Buhling et al., 2011; Thorenoor et al., 2018)), markedly increased with disease severity. Both SFTPA2 and CTSH are synthesized in type II pneumocytes (Buhling et al., 2011; Thorenoor et al., 2018), which are primary targets for SARS-CoV-2 infiltration. As such, these proteins may be early markers of type II pneumocyte dysfunction, impaired surfactant synthesis and lung damage as SFTPA2 plasma levels have been shown to be elevated in patients suffering from acute respiratory failure (Doyle et al., 1997). These results are consistent with recent transcriptomics analyses of lung biopsies showing impaired surfactant production in patients with severe COVID-19 (Islam and Khan, 2020).

Key proteomic features of the COVID-19 severity model also included two components of the RAS (ACE2 and MME), a complex hormonal system that regulates blood pressure, fluid homeostasis as well as pulmonary inflammation (Chappell, 2019). The involvement of the RAS in the pathogenesis of SARS-CoV-2 infection is well-established since the interaction between the viral Spike protein and ACE2 is a primary mechanism of viral entry into host cells (Mortaz et al., 2020). High ACE2 and MME plasma levels in severe patients suggest increased shedding from the cell surface which may result in loss of function of these proteins (Ni et al., 2020). A loss of function of ACE2 and MME will likely result in a decrease of angiotensin I (Ang-I) and angiotensin II (Ang II) degradation, which aligns with the multi-organ injuries observed in severe patients (Liu et al., 2020; Ni et al., 2020). Interestingly, increased plasma ACE2 concentration has recently been associated with an increased risk for subsequent cardiovascular events in patients with COVID-19 (Narula et al., 2020). As such, our analysis points at mechanistic markers of disease severity that may also be implicated in the clinical manifestations of patients recovering from COVID-19, such as cardiovascular or neurological complications.

The most informative protein feature in our severity model was MERTK, a membrane-bound receptor tyrosine kinase that plays a role in immunoregulation, hemostasis, and endothelial barrier integrity (Cai et al., 2018; Lemke and Silverman, 2020; Li et al., 2019), whose levels in plasma substantially increased with severity. Loss of endothelial MERTK expression in the lung can result in enhanced permeability and leukocyte transendothelial migration, thereby aggravating inflammation at the site of infection and contributing to acute respiratory distress syndrome (Li et al., 2019). In addition, soluble MERTK can hinder activation of membrane-bound MERTK, resulting in the inhibition of macrophage-mediated clearance of apoptotic cells and the loss of its immunosuppressive effect (Cai et al., 2018; Lemke and Silverman, 2020; Sather et al., 2007). Through its role in both endothelial integrity in the lung and immunoregulation, dysregulated MERTK could potentially be a central player in the development of severe COVID-19. In addition, MERTK also plays a role in platelet aggregation (Sather et al., 2007), and its observed higher plasma levels in patients with the severe disease together with increased plasma levels of other proteins involved in hemostasis (Hrycek and Cieslik, 2012; Ikeda et al., 2018; Lu et al., 2020a) suggests an increased risk for coagulation and thrombosis. Coagulopathy and thrombosis have been observed in many patients that presented with severe COVID-19 (Gao et al., 2020; Levi et al., 2020).

This study has certain limitations. First, in this cohort, we only assessed peripheral blood samples of patients affected by COVID-19. It would be of great interest to assess the local immune and proteome perturbations in parallel by analyzing lung biopsies or bronchoalveolar lavage fluid to investigate whether the same trends and dampened immune cell responses are observed locally in the lung as well. Second, this is a single-center study with samples collected from patients at Stanford University Medical Center. Most of the patients that were encountered during the study period had mild symptoms. Future studies including patients with different ethnicity, socio-economic status, and co-morbidities will be important to test the boundary of generalizability of our findings. Similarly, we recognize that the severity model is independently validated as a high-dimensional model. Additional testing in larger and more diverse patient cohorts is necessary to determine whether individual model features can serve as clinical biomarkers of disease severity. Finally, this is a cross-sectional study --longitudinal monitoring of plasma and single-cell proteomic features will give increased insight into the fluctuation of these features over time and lead to new insights into patients with COVID-19 “long-hauler” status.

Determining the underlying immune pathogenesis across the spectrum of COVID-19 severity remains an important clinical challenge. Our integrated analysis of plasma and single-cell proteomics in patients with mild, moderate, and severe COVID-19 identified a multivariate model that differentiates COVID-19 severity. The observations identified by this model contribute clinically relevant insights into the status of patient’s immune responses during SARS-CoV-2 infection and provide promising severity-specific biological markers for future validation that may inform decision-making on potential therapeutic targets for the prevention of disease progression.

## Materials and methods

### Study design and sample collection

#### Design

We conducted this study at Stanford University Medical Center, where the samples were collected at the Stanford Occupational Health Clinic under an IRB approved protocol (55689; Protocol Director Dr. Nadeau). We obtained samples from adults with positive test results for SARS-CoV-2 from analysis of nasopharyngeal swab specimens obtained at any point from March to June 2020. Follow-up continued through October 5, 2020. Testing was accomplished using Stanford Health Care clinical laboratory developed internal testing capability with a quantitative reverse-transcriptase–polymerase-chain-reaction (qRT-PCR) assay. Informed consent was obtained from each patient prior to enrolling in the study or from the patient’s legally authorized representative if the patient was unable to provide consent. Participants were excluded if they were taking any experimental medications (i.e. those medications not approved by a regulatory agency for use in COVID-19). Healthy controls were collected prior to the detection of SARS-CoV-2 in the region (historical controls) and were consented using a separate IRB-approved protocol (Protocol Director Dr. Nadeau).

#### Data sources and clinical definitions

We obtained data from self-reported surveys and from Stanford clinical data electronic medical record system as per consented participant permission. This database contains all the clinical data available on all inpatient and outpatient visits to Stanford facilities. The data obtained included patients’ demographic details, vital signs, laboratory test results, medication administration data, historical and current medication lists, historical and current diagnoses, clinical notes, radiological results, biopsy results as appropriate, historical discharge disposition for previous inpatient hospitalizations, and ventilator use data. Patient data were collected and definitions and diagnosis of disease were used according to previously defined criteria (Chen et al., 2020) with the clinical classification of COVID-19 as followed; Mild disease: mild to no symptoms, and no manifestation of pneumonia; Moderate disease: fever and respiratory tract symptoms; Severe disease: hospitalization with respiratory distress.

#### Phlebotomy and initial blood processing

Blood was analyzed from samples collected from 97 patients with COVID-19 and 40 controls via venipuncture. Anticoagulated blood was processed into peripheral blood mononuclear cells (PBMC) by density gradient centrifugation using published methods (Syed et al., 2014). PBMC were stored in 10% DMSO and frozen in liquid nitrogen until thawing and staining. Plasma was obtained from blood collected in EDTA tubes using standard procedures and stored at −80°C.

### Mass cytometry analysis of single-cell immune cell responses in PBMC

#### In vitro PBMC stimulation

Cryopreserved PBMC were quickly thawed, washed two times with supplemented medium, and rested for 1h at 37°C in RPMI 1640 medium supplemented with 10% fetal bovine serum, 1% Penicillin-Streptomycin, and 1% L-Glutamine. PBMC were counted and checked for viability. 0.5-1×10^6^ cells were either stimulated with lipopolysaccharide (LPS; 1*μ*g/ml) and CL097 (TLR7/8 agonist; 1*μ*g/ml), interleukin-2 (IL-2), IL-4, IL-6 and interferon-*α* (IFN*α*; all 100 ng/ml), a cocktail of phorbol 12-myristate 13-acetate (PMA), ionomycin, brefeldin A and monensin (1x; PI cocktail), or left unstimulated for 15 minutes at 37°C. After stimulation, samples were fixed with Proteomic Stabilizer (SmartTube) and stored at −80°C until further processing for mass cytometry analysis.

#### Barcoding and antibody staining

The 42-marker mass cytometry antibody panel included 25 cell surface antibodies and 17 intracellular antibodies recognizing primarily phospho-specific signaling epitopes (**Supplemental data file S2**). In brief, following *in vitro* stimulation, fixed PBMCs were thawed, reconstituted in cell staining medium, and arranged in a 96-well block. Subsequent steps were performed using a previously described robotics platform (Bjornson-Hooper et al., 2019). Sets of 16 samples were barcoded with palladium metal (Zunder et al., 2015) and pooled into a single well. Pooled barcoded samples were treated with FC-block (Human TruStain FcX, Biolegend) for 10 minutes then surface antibody stained for 30 minutes in cell staining medium (PBS with 0.5% BSA and 0.02% sodium azide). After surface staining, cells were permeabilized in ice-cold 100% methanol, washed, and stained for 60 minutes with intracellular antibodies in cell staining medium. Following intracellular staining, cells were washed and resuspended in an iridium intercalator (Fluidigm) solution containing 1.6% paraformaldehyde. Finally, samples were washed, resuspended in 1X five-element normalization beads (La, Pr, Tb, Tm, Lu) (Fluidigm), and analyzed on a freshly cleaned and tuned CyTOF instrument. The resulting mass cytometry data were bead-normalized across all runs and debarcoded as previously described (Finck et al., 2013).

#### Cell frequency, baseline intracellular signaling, and intracellular signaling responses

Mass cytometry data was examined using CellEngine (Primity Bio) to define cell populations using manual gates and quantify differential expression of signaling markers in response to stimulation. The gating strategy can be found in **supplemental figure S1**. Cell frequencies were expressed as a percentage of gated singlet mononuclear cells (DNA^+^CD235a^-^CD61^-^CD66^-^), except for granulocyte frequency which was expressed as a percentage of singlet leukocytes (DNA^+^CD235a^-^CD61^-^). Phospho-signal intensity was quantified per single cell for each signaling protein (pSTAT1, pSTAT3, pSTAT4, pSTAT5, pSTAT6, pMAPKAPK2, pCREB, pPLC_γ_2, pS6, pERK1/2, pP38, pZAP70/Syk, pTBK1, p4EBP1, and total IkB*α*) using an arcsinh transformed value (arcsinh(x/5)) from the median signal intensity. Endogenous or baseline intracellular signaling activity was derived from the analysis of unstimulated cells, while intracellular signaling responses to stimulation were reported as the arcsinh transformed ratio over the baseline signaling, i.e., the difference in arcsinh transformed signal intensity between the stimulated and unstimulated condition. For cell subsets in a given sample that had an event count below 20 events, that cell subset and related phospho-signal were excluded from downstream analysis. A penalization matrix, based on mechanistic immunological knowledge, was applied to the immune cell response data (Culos et al., 2020; Ghaemi et al., 2019).

### Plasma Protein Profiling Using Olink Multiplex Panel

Plasma protein levels were quantified using Olink multiplex proximity extension assay (PEA) panels (Olink Proteomics; www.olink.com) according to the manufacturer’s instructions and as described before (Assarsson et al., 2014). The basis of PEA is a dual-recognition immunoassay, where two matched antibodies labeled with unique DNA oligonucleotides simultaneously bind to a target protein in solution. This brings the two antibodies into proximity, allowing their DNA oligonucleotides to hybridize, serving as a template for a DNA polymerase-dependent extension step. This double-stranded DNA which is unique for a specific antigen will get amplified using P5/P7 Illumina adapters along with sample indexing, which is quantitatively proportional to the initial concentration of target protein. These amplified targets will finally get quantified by Next Generation Sequencing using Illumina Nova Seq 6000 (Illumina Corporation. San Diego, California). In this study, we have used the Explore 1536 panel which measures 1,472 proteins using 3ul plasma samples, which were treated with 1% Triton X-100 and incubated at room temperature for 2 hours to inactivate the virus. The raw expression values obtained with the Olink assay are provided in the arbitrary unit Normalized Protein Expression (NPX), where high NPX values represent high protein concentration. Values were log2-transformed to account for heteroskedasticity. Proteins close to the limit of detection are flagged in the raw data.

### Statistical analysis

#### Univariate analysis

We used the R environment (http://www.r-project.org/) for statistical analysis. We chose to apply a ranked regression analysis for each feature relative to the severity of the patient at the time of sampling. Multiple comparison corrections were performed using the Benjamini-Hochberg procedure (Benjamini and Hochberg, 1995). For each statistic, we report both p-values, the associated FDR corrected “q-values,” and regression estimates.

#### Multi-omic factor analysis

Multi-omic factor analysis (MOFA) was applied simultaneously across plasma (Olink) and the single-cell proteomics data (cell frequency, baseline signaling, IFN*α*/IL-2/IL-4/IL-6 signaling response, LPS/CL097 signaling response, and PI signaling response). MOFA infers a set of factors that capture both biological and technical sources of variability that are shared across different datasets. MOFA models were constructed using the six data layers which were supplied as a list of matrices. We followed the developers’ directions for model selection and downstream analysis (Argelaguet et al., 2018). Since MOFA is not guaranteed to find a global optimum, 10 model trials were performed using different random initializations. For each trial, the number of factors was calculated by requiring at least 2% variance explained for any single dataset. The model with the highest evidence lower bound was selected out of these 10 trials (model 6, **Figure S2A**). The factors calculated by model 6 of our 10 trials were extracted for downstream analysis. MOFA enables variance decomposition of the calculated factors and uses a coefficient of determination (R^2^) to quantify the fraction of variance explained by each factor for each dataset, which we examined first to determine how each dataset contributes to each factor (**Figure S2B**). Next, we regressed the 17 factors on COVID-19 severity, which was dummy encoded as follows: Control = 1, Mild = 2, Moderate = 3, Severe = 4. Regression estimates, 95% confidence intervals, and p-values were examined (**Figure S2C**). Factors 3, 8, and 10 were significantly associated with COVID-19 severity. To assess the robustness of factors across model trials, we calculated Pearson correlation coefficients between every pair of factors across all trials. Factors 3 and 10 were consistently discovered in all model instances (data not shown). Finally, we visualized our samples across factors 3 and 10 using a bivariate scatterplot (**Figure 1D**).

#### Multivariate analysis and stacked generalization

For the multivariate analysis, a LASSO model was trained independently on each omics dataset independently. For a matrix X of all biological features from a given -omic data set, and a vector of disease severity Y, the LASSO algorithm (Tibshirani, 1996) calculates coefficients β to minimize the error term L(β)=||Y−Xβ||2. The L1 regularization is used to increase model sparsity for the sake of biological interpretation and model robustness. Once a LASSO model is trained for each omics modality (individual model performance **Table S2**), the multi-omics analysis can be carried out by performing stacked generalization (Ghaemi et al., 2019) on the new representation of the data by using the outputs of the previous layer of models as predictors with the constraint that their participation should be a non-negative coefficient. Particularly, a LASSO model is first constructed on each omics modality, then all estimations of disease severity are used as predictors for a second-layer constrained regression model. Intrinsically, this is equivalent to a weighted average of the individual models with the coefficients of the LASSO model as desired weights.

#### Cross-validation

An underlying assumption of the LASSO algorithm is statistical independence between all observations. In this analysis, subjects are independent, and, in this regime, a model is trained on all available samples from all subjects. However, at each iteration, one sample is kept for independent validation. The reported results are exclusively based on the blinded subject. For stacked generalization, a two-layer cross-validation strategy was implemented where the inner layer selects the best values of λ and reports the intermediate prediction. Then, the outer layer optimizes the constrained regression of all predictors for the stacked generalization step. Cross-validation folds are carefully synchronized between the individual models from each omics.

#### Bootstrap analysis for feature selection

For each omics dataset, we performed a bootstrap analysis where we repeat a bootstrapping procedure on the dataset and train a cross-validated model. At each iteration, we keep the non-zero coefficients selected by the LASSO procedure on the bootstrapped dataset and we repeat the procedure 1,000 times (Bach, 2008; Chatterjee and Lahiri, 2011). We report the frequency of selection of the features as well as their median coefficient in all the bootstrap models. This method selected 44 frequency, 599 baseline, 536 PI-response, 492 IFN*α*/IL-2/IL-4/IL-6, and 783 proteome features that were informative in at least one iteration of the bootstrap procedure (see **supplemental file S1**). To assess the relative importance of each feature to the model, we ranked features in each data layer based on their frequency of selection.

#### Correlation network

The features are visualized using a correlation graph structure to identify correlated feature populations. Each biological feature is denoted by a node whose size is dependent on the *-log_10_* of p-value of correlation with disease severity (Spearman). The correlation network between the features is represented by an edge where the width of the edges is proportional to the Spearman p-value of the correlation between a pair of nodes on a log 10 scale. The graph is visualized using the t-SNE layout.

#### Confounder analysis

A linear regression analysis was used as a statistical method to exclude the likelihood that certain clinical or demographic variables confounded the predictive accuracy of the severity model.

#### Multi-class receiver operating characteristics (AUC)

To characterize our predictions’ separability in a multi-class setting, we used a combined metric of the area under the receiver operating curve (AUC) in both the training and validation model. This metric for multi-class analysis uses every combination of labels in one-to-one comparisons. Every subset of predictions on a pair of classes is considered independently, and the AUC is calculated for each specific pair. Once all the subset AUCs are computed, the generalizable AUC combines them, taking their mean across all comparisons, to give the multi-class model’s overall performance (Hand and Till, 2001).

#### Plasma proteomic pathway identifier

To obtain pathway information of selected proteome features, we utilized Reactome (www.reactome.org), a web-based resource for identifying biological pathways, in which we used the list of the 10% bootstrap-selected proteins as an input. Reactome provided a list of pathways, identified by a Reactome gene set identifier, which we assessed for the proteins part of specific pathways.

## Supporting information

Antibody panel

Model features selected by bootstrap procedure

## Data availability

Raw data, processed data, and source code for the reproduction of the results will be made publicly available on Flowrepository (ID: FR-FCM-Z3G5) and Github.

## Supplemental material

Supplemental file S1 lists the features informative of the severity model as selected by the bootstrap procedure (xlsx-file). Supplemental file S2 contains the mass cytometry antibody panel. Figure S1 shows the manual gating strategy used to identify innate and adaptive immune cells. Figure S2 shows the MOFA analysis to identify shared sources of variability across the plasma and single-cell proteomics data. Figure S3 displays the frequency of plasmablasts, and the immune cell signaling features at baseline and after stimulation with inflammatory agents that are informative to the severity model. Figure S4 shows plasma protein level features that are informative to the severity model. Figure S5 shows plasma protein level features that are part of neutrophil degranulation and hemostasis, and informative to the severity model. Table S1 contains the model performance of the individual data layers, and the contribution of the individual data layers to the severity model (weighted average of model coefficients). Table S2 shows how well the severity classifies each patient group from the other. Table S3 contains the results of the confounder analysis.

## Author contributions

Conceptualization: BG, KCN, DRM, GN

Data curation: KCN, GKRD, JF, MM, JH, DF

Formal Analysis: JH, JG, DF

Funding acquisition: BG, KCN, DRM, GN

Investigation/Data acquisition: DF, HC, EST, DRM, LSP, KA, MM, ED, GKRD, JF, IC, TTS, CW, JM, EAG, IAS, XH, FV, DKG, AST, KKR, SJ, SIVF, DJK, DaF

Methodology: BG, DRM, GPN, NA, JH Project administration: BG, DRM, KCN Resources: KCN, DRM, BG, NA, BAP, SDB, MSA, MA, SC, RW Software: NA, JH, JG

Supervision: BG, DRM, KCN, GPN Visualization: DF, JH, HC, JG, AST Writing – original draft: BG, DRM, DF

Writing – review & editing: BG, DRM, KCN, DF, JH, JG, NA and all authors

## Acknowledgments

This work was supported in part by the U.S. Food and Drug Administration Medical Countermeasures Initiative contracts 75F40120C00176 and HHSF223201610018C (GPN, DRM, SJ, HC); the Bill and Melinda Gates (BMGF) foundation INV-002704 (GPN), OPP1113682 (GPN), COVID-19 Pilot Award OPP1113682 (GPN, DRM, SJ, HC), Center for Human Systems Immunology Pilot Seed grant (BG); the Doris Duke Charitable Foundation (2018100A, BG); a CEND Covid Catalyst Award (BG); the Mercatus Center Fast Grants for COVID-19 (BG, GPN, DRM, SJ, HC); the National Institutes of Health R35GM137936 (BG), SeroNet 1U54CA260517 (SDB and KCN), CCHI U19AI057229 (KCN and SDB), K23 grant number AI135037 (JDK), T32GM089626 (JGM), R35GM138353 (NA), 5U19AI100627-09 Systems Approach to Immunity and Inflammation COVID supplement, National Cancer Institute Task Order No. HHSN261100039 under Contract No. HHSN261201500003I; Burroughs WellCome Fund (NA); the Sean N Parker Center for Allergy and Asthma Research (KCN, RSC); the Sean N Parker Institute for Cancer Immunotherapy (G.P.N.); the Sunshine Foundation (KCN, RSC); the Crown Foundation (KCN, RSC); the Centers for Disease Prevention and Control contract number 75D30120C08009 (JDK); the Consejo Nacional de Ciencia y Tecnología (CONACyT, Mexico) grants 289788 and 311783 (SIVF); and the Rachford and Carlota A. Harris Endowed Professorship (GPN). This article reflects the views of the authors and should not be construed as representing the views or policies of the FDA, CDC, NIH, or other funding sources listed here. The authors declare no competing financial interests.

## Supplementary materials for

**Figure S1.**
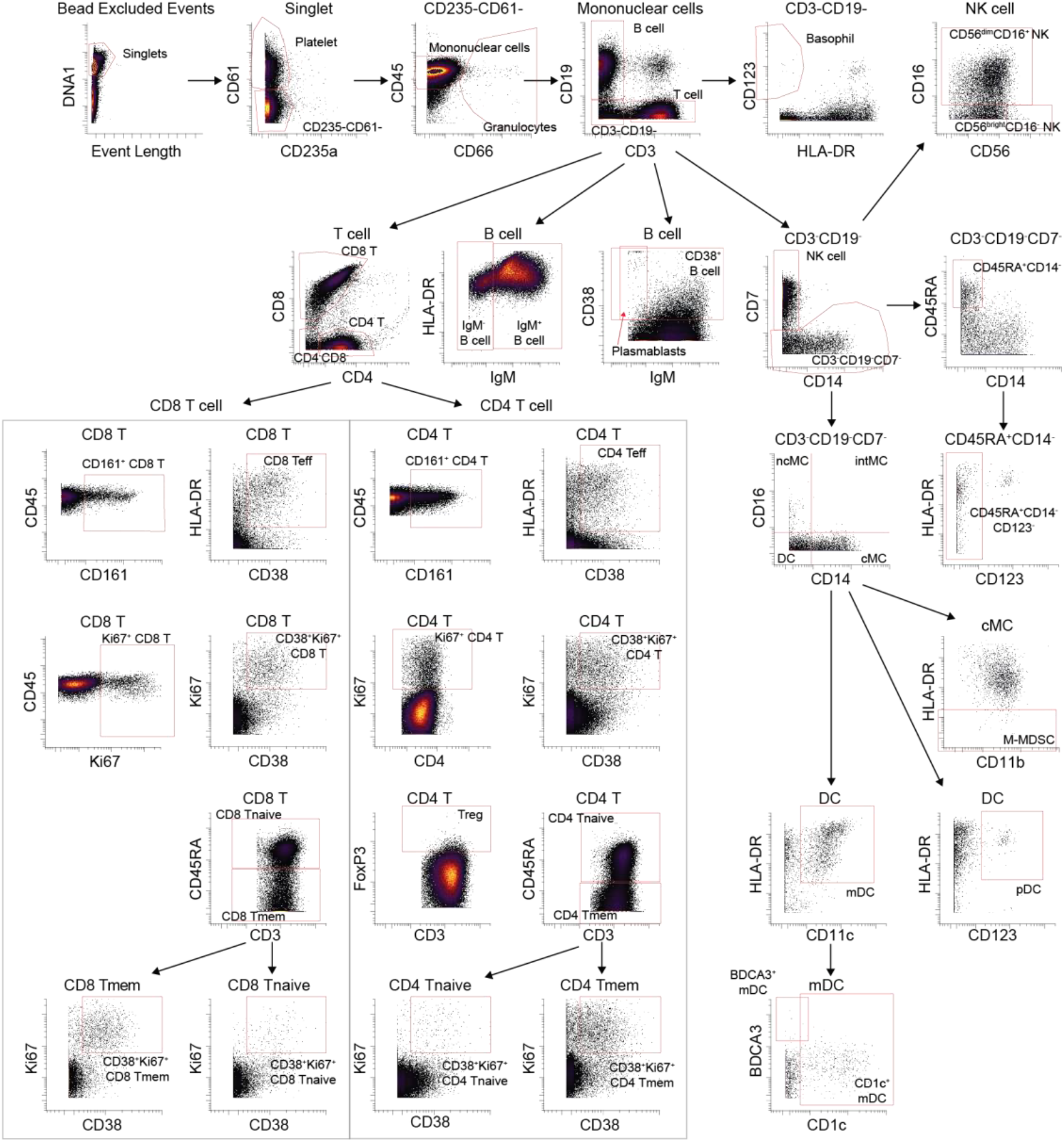
Hierarchical manual gating strategy used for mass cytometry analyses.

**Figure S2.**
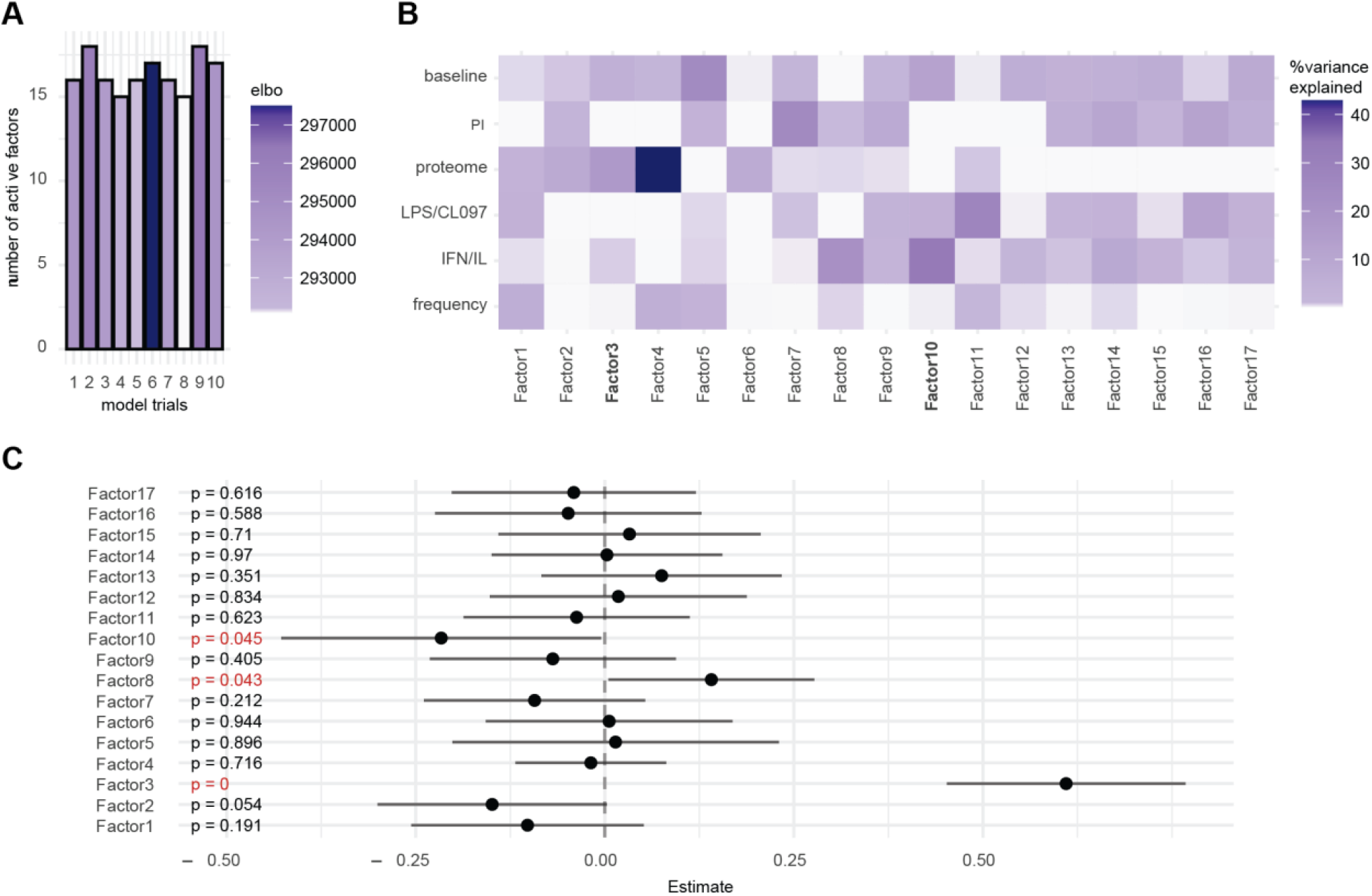
Multi-omic factor analysis (MOFA) identifies shared sources of variability across single-cell cytometry and proteomic data that are associated with COVID-19 severity. **A** Ten MOFA models were constructed with different random initializations. Shown are the number of factors calculated for each trial, and colored by the corresponding evidence lower bound (ELBO). **B** Portion of variance explained (R^2^) for each factor and for each dataset. IFN/IL = IFN*α*/IL-2/IL-4/IL-6 **C** Association of MOFA factors with COVID-19 severity (encoded as Control = 1, Mild = 2, Moderate = 3, Severe = 4) was performed with linear regression analysis. Estimates, 95% confidence intervals, and p-values are shown (significant p-values highlighted red).

**Figure S3.**
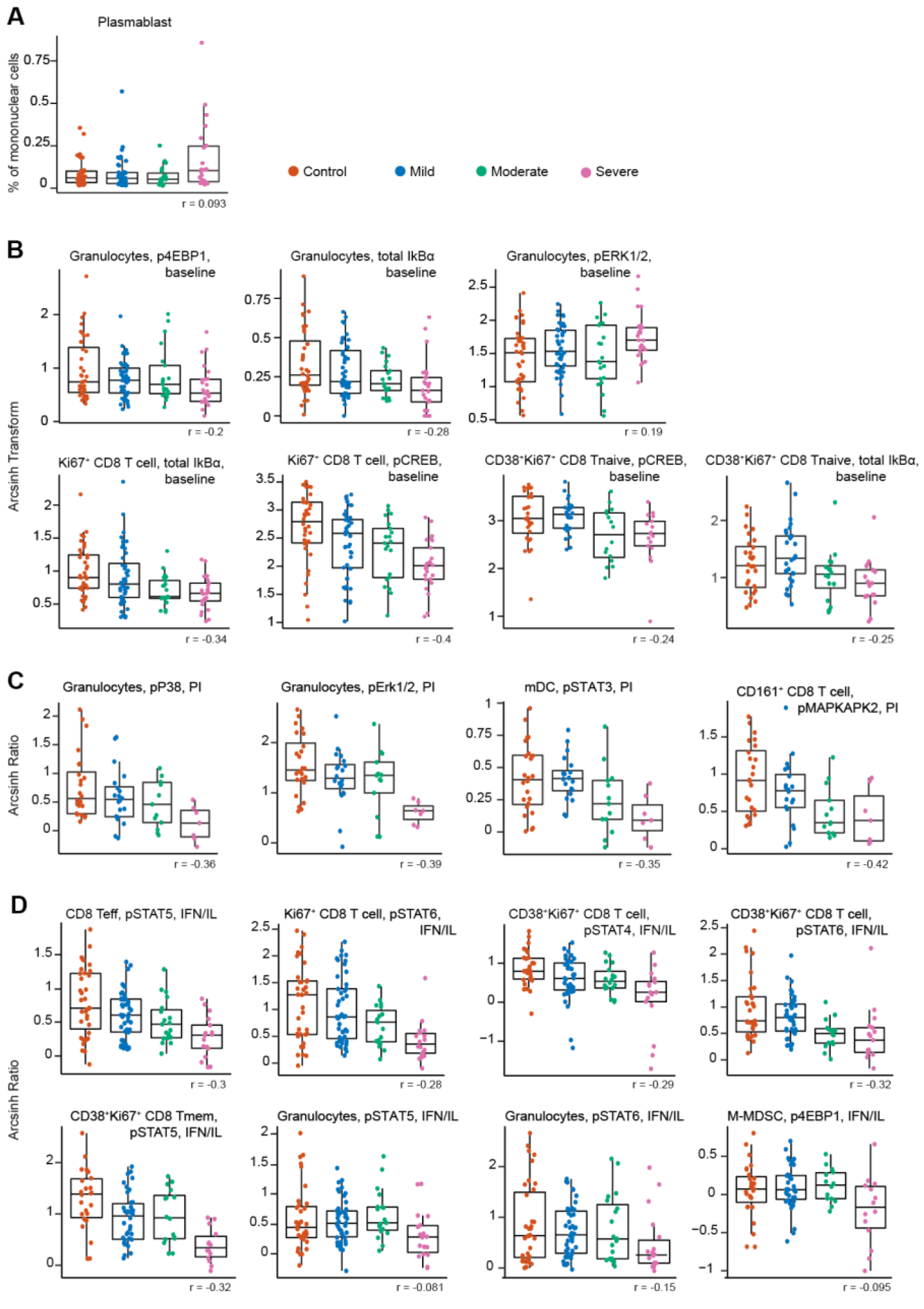
Frequency and immune cell signaling response features. Boxplots showing plasmablast frequency, immune cell signaling responses at baseline (reported as arcsinh transformed values), and signaling response after stimulation with inflammatory agents (reported as arcsinh transformed ratio over the baseline signaling response; see methods). **A** Plasmablast frequency (CD38^+^IgM^-^CD19^+^). **B** Baseline immune cell signaling response. **C** immune cell signaling response to PI stimulation, and **D** response to IFN*α*/IL-2/IL-4/IL-6 stimulation (IFN/IL) of features informative in the COVID-19 severity model. Boxplots classified by disease severity and including correlation of the feature with disease severity (Spearman coefficient). Tmem = memory T cell; Teff = effector T cell.

**Figure S4.**
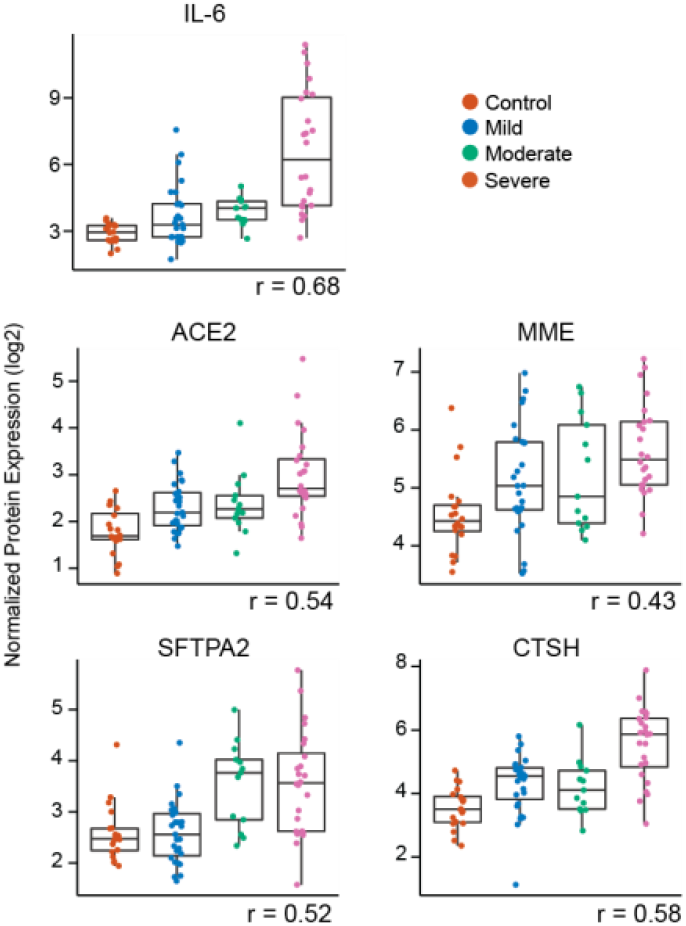
Plasma proteomic features informative to the model. Boxplots showing plasma protein levels of features that are informative in the COVID-19 severity model. Boxplots are classified by disease severity and include the correlation of the feature with disease severity (Spearman coefficient). ACE2 = angiotensin-converting enzyme 2; MME = neprilysin; SFTPA2 = surfactant-associated protein A2; CTSH = cathepsin H.

**Figure S5.**
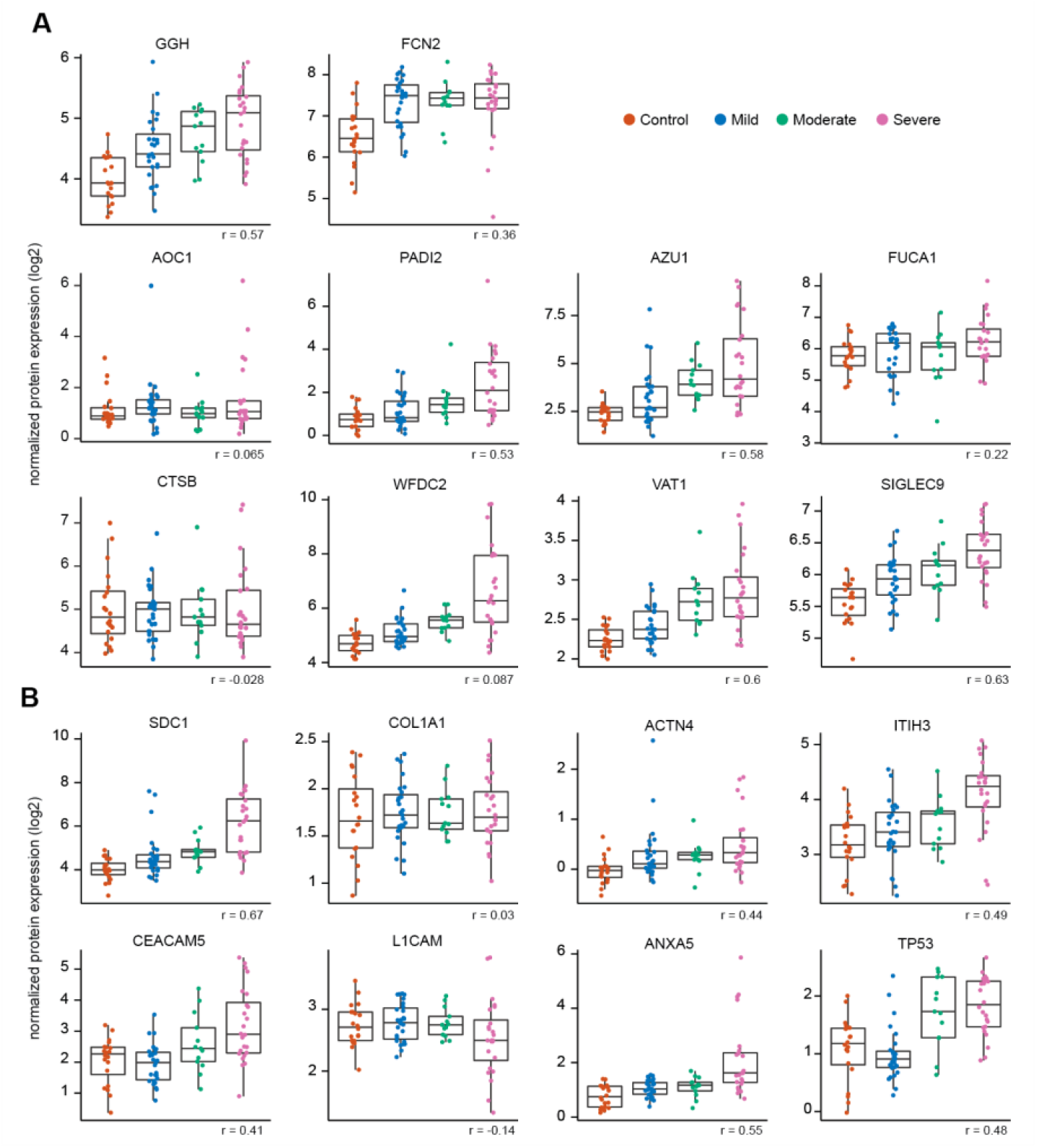
Informative plasma proteomic features involved in neutrophil degranulation and hemostasis. Boxplots showing plasma protein levels of features that are informative in the COVID-19 severity model and that are part of **A** neutrophil degranulation (Reactome gene set identifier R-HSA-6798695.2) and **B** hemostasis (Reactome gene set identifier R-HSA-109582). Boxplots are classified by disease severity and include the correlation of the feature with disease severity (Spearman coefficient). GGH = Gamma-glutamyl hydrolase; FCN2 = Ficolin-2; AZU1 = Azurocidin; FUCA1 = Tissue alpha-L-fucosidase; AOC1 = Amiloride-sensitive amine oxidase [copper-containing]; PADI2 = Protein-arginine deiminase type-2; VAT1 = Synaptic vesicle membrane protein VAT-1 homolog; SIGLEC9 = Sialic acid-binding Ig-like lectin 9; CTSB = Cathepsin B; WFDC2 = WAP four-disulfide core domain protein 2; SDC1 = syndecan-1; COL1A1 = Collagen alpha-1 (I) chain; ACTN4 = Alpha-actinin-4; ITIH3 = Inter-alpha-trypsin inhibitor heavy chain H3; CEACAM5 = Carcinoembryonic antigen-related cell adhesion molecule 5; L1CAM = Neural cell adhesion molecule L1; ANXA5 = Annexin A5; TP53 = Cellular tumor antigen p53.

**Table S1.**
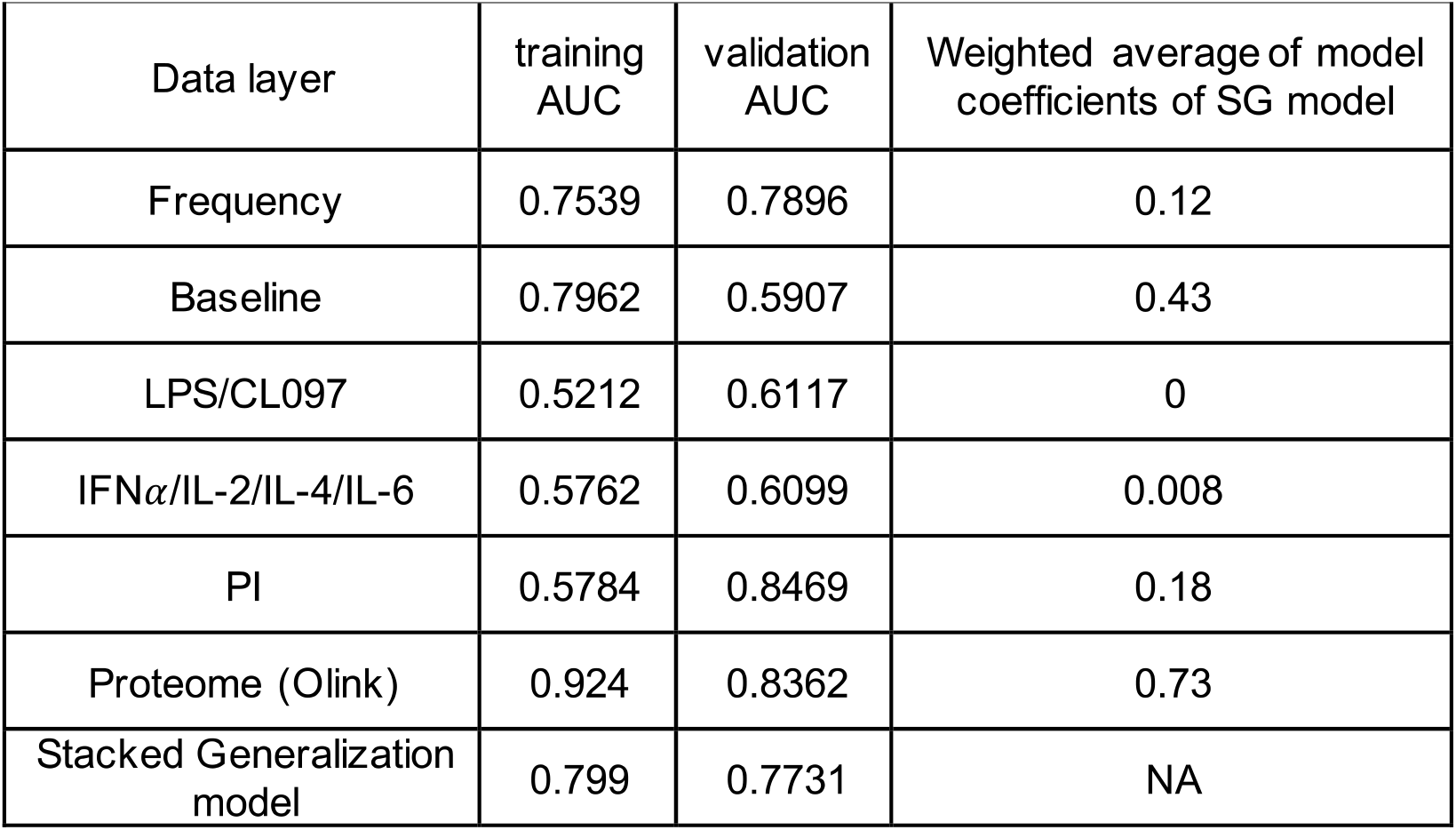
Model performance and weighted average of individual data layers. AUC indicates the model performance of individual data layers in classifying severity. Weighted average shows the contribution of individual data layers to the severity model. Results related to Figure 2 B-C. SG model = Stacked Generalization model

**Table S2.** AUC of individual ROC analyses for performance of severity model to classify severity. AUC shows how well the severity model is able to classify each patient group from the other. 1 = control, 2 = mild, 3 = moderate, and 4 = severe. Results related to Figure 2 D-E

|  | training AUC | validation AUC |
| --- | --- | --- |
| 1vs2 | 0.71 | 0.6071 |
| 1vs3 | 0.85 | 0.7697 |
| 1vs4 | 0.90 | 0.9238 |
| 2vs3 | 0.71 | 0.6623 |
| 2vs4 | 0.82 | 0.8571 |
| 3vs4 | 0.81 | 0.8182 |

**Table S3.** Confounder analysis. None of the potentially confounding variables significantly influenced the COVID-19 severity model. Related to Figure 2. SexM = male sex. BMI = body mass index. NS = not significant

| Variable | Estimate | Std.Error | t | value | Pr(>\|t\|) |
| --- | --- | --- | --- | --- | --- |
| Prediction model | 0.441365 | 0.128161 | 3.444 | 0.00177 | *** |
| Days between symptom onset and sample collection | -0.005135 | 0.006448 | -0.796 | 0.43225 | NS |
| Age | 0.010871 | 0.006316 | 1.721 | 0.09589 | NS |
| SexM | -0.306411 | 0.199897 | -1.533 | 0.13615 | NS |
| BMI | 0.050989 | 0.016994 | 3.000 | 0.00550 | ** |

